# The genomic architecture of a continuous color polymorphism in the European barn owl (*Tyto alba*)

**DOI:** 10.1101/2023.04.03.535036

**Authors:** Tristan Cumer, Ana Paula Machado, Luis M. San-Jose, Anne-Lyse Ducrest, Céline Simon, Alexandre Roulin, Jérôme Goudet

**Affiliations:** Department of Ecology and Evolution, University of Lausanne, Lausanne, Switzerland; Laboratoire Évolution and Diversité Biologique, UMR 5174, CNRS, Université Toulouse III Paul Sabatier, Toulouse, France; Swiss Institute of Bioinformatics, Lausanne, Switzerland

**Keywords:** QTL mapping, Color polymorphism, Selection, Melanin

## Abstract

The maintenance of color polymorphism in populations has fascinated evolutionary biologists for decades. Studies of color variation in wild populations often focus on discrete color traits exhibiting simple inheritance patterns, while studies on continuously varying traits remain rare. Here, we studied the continuous white to rufous color polymorphism in the European barn owl (*Tyto alba*). Using a Genome Wide Association approach on whole-genome data of 75 barn owls sampled across Europe, we identified, in addition to a previously known *MC1R* mutation, two regions involved in this color polymorphism. We show that the combination of the three explains 80.37% (95% credible interval 58.45 to 100%) of the color variation. Among the two newly identified regions, the one on the sexual chromosome (Z) shows a large signal of differentiation in the Swiss population when contrasting individuals with different morph but the same *MC1R* genotype. We suggest it may play a role in the sexual dimorphism observed locally in the species. These results, uncovering two new genomic regions, provide keys to better understand the molecular bases of the color polymorphism as well as the mechanisms responsible for its maintenance in the European barn owl at both continental and local scales.

## Introduction

Color polymorphism, and the mechanisms that allow its maintenance in populations, have fascinated evolutionary biologist for decades. Pioneer studies in wild populations, such as Kettlewell’s studies [1] on the action of natural selection on the black and pale morphs of the peppered moth, *Biston betularia*, or Endler’s studies [2] on the joint role of sexual and natural selection on explaining color variation in guppies, *Poecilia reticulata*, have shaped our understanding of the evolution and maintenance of color polymorphism. For practical reasons, research on animal coloration has continued and flourished mainly on humans [3–5] and model systems (i.e. mice [6,7], or domestic animals [8,9]), and has been more recently extended to non-model species, with a handful of studies on birds [10,11], mammals [12,13], butterflies [14] and amphibians [15]. Moreover, many of these studies took advantage of color traits exhibiting relatively simple discrete variation and inheritance patterns thus often contrasting clearly distinctive (eco)morphs [11-15]. It is however known that many color traits vary continuously between two extreme values (for instance, human skin color [16]), and that the expression of these traits can be determined by genetic factors (for example, color in the arctic Fox [13]), by the environment (for instance the flamingo color acquired through their food [17]) or by the interaction of genetic and environmental factors (such as human skin color, determined by genetics [3–5] and exposition to UVs [16]). Thus, our current understanding of the genetics basis of animal coloration is limited to a handful of well-studied systems, and rarely accounts for the continuous character of the color variation. Further work is therefore needed to better characterize the molecular basis of the color diversity observed in many species.

Barn owls (Tytonidae) represent an ideal system to study continuous color variation between individuals and populations. These cosmopolitan birds represent an extraordinary example of phenotypic variation, with plumage color clines across continents found in at least seven *Tyto* clades [18]. Among them, the color cline in Europe is the most pronounced, with individuals ranging from white in the south to dark rufous in the north of its distribution [19]. Previous studies have shown that this melanin-based coloration is associated with reproductive success and feeding rate [20], habitat choice [21], diet [22] wing morphology and stomach content while flying [23,24], as well as dispersal ability [25,26]. The association between the color and a wide variety of traits may be induced by the strong pleiotropy of the melanocortin system controlling numerous traits [27], among which the production of eu/pheo-melanin pigments responsible for the coloration of the barn owl [28,29]. The rufous color variation of the barn owl is known to be strongly heritable (h^2^ of the ventral body side color of owls ranging from 0.57 to 0.84; [30]). A diallelic mutation (*V126I*) in the melanocortin-1 receptor gene (*MC1R*) explains a large proportion of the phenotypic variance in reddish coloration [31], with individuals homozygous for the *MC1R* allele with a valine at position 126 (allele denoted *MC1R-white*, genotype *MC1R*_*VV*_ below) being whitish to light rufous, and individuals with at least one isoleucine being rufous (allele denoted *MC1R-rufous*, genotypes *MC1R*_*VI*_ and *MC1R*_*II*_ below). Interestingly, a recent study reconstructing the demographic and colonization history of the barn owl in Europe showed that the color cline is not specific to a lineage [32], de-coupling the color and the neutral history of the populations.

Considering the association between the amount of rufous coloration and several biotic and abiotic factors, combined with its unknown evolutionary origin, the question of the mechanisms underlying the maintenance of this polymorphism remains elusive. So far, three non-mutually exclusive mechanisms have been proposed. First, color itself may be a primary target of local adaptation at continental scale, favoring the rufous form in the north and whitish form in the south. We [19,33,34] previously hypothesized that the clinal variation in coloration may be maintained by natural selection, since phenotypic differentiation between populations across the European cline is much more pronounced that neutral genetic differentiation. A role of foraging has been suggested in this context [22], possibly because a whitish plumage reflects moonlight which induces fear in their prey [35]. This local adaptation hypothesis is further supported by the higher frequency of the isoleucine *MC1R* variant in northern population while it is nearly absent in southern populations [19]. Second, at local scale, density-dependent selection on the different morphs may maintain the polymorphism in the populations. Kvalnes et al. [31] pointed that rufous females were selected for at low densities, while whitish ones were favored at high densities. Thus, the fluctuation of population density could cause the selection for rufous or whitish form, allowing for the maintenance of the polymorphism. Finally, a sexually antagonist selection may also be acting in the populations: a recent study pointed that dark melanic females (i.e., harboring a rufous plumage with many black spots located at the tip of ventral body feathers) were sexually mature earlier than lighter melanic females while lighter melanic males (i.e., harboring a whitish plumage with few black spots located at the tip of ventral body feathers) were sexually mature earlier than darker melanic males [36]. Combined with the fact that individuals that matured faster produced a larger number of fledglings per year than individuals that matured slower [36], these results suggested that a light melanic plumage is beneficial in males and a dark melanic plumage in females and indicate that sexually antagonist selection may be at play in maintaining this polymorphism. Finally, a potential epistatic effect of the *MC1R-rufous*, masking the expression of other variants responsible of the variation of color between individuals carrying the *MC1R-white* allele, may also play a role in maintaining the polymorphism by hiding some variants from selection [37].

Thus, conclusive evidence for the selective targets and agents establishing the European barn owl color polymorphism and cline is still amiss. Identifying the gene(s) underlying barn owl plumage coloration and their effects is a first step to unravel the molecular basis of this polymorphism.

In the present study, we investigated the genomic basis of color polymorphism in the European barn owl. By exploiting whole genome data of 75 barn owl individuals (*T. alba*) from 6 different populations / localities from Europe and the Middle East, combined with spectrophotometric data on their coloration; we (i) identified major Quantitative Trait Loci (QTL), (ii) studied the levels of variation explained by these QTL variants, (iii) discuss the potential functional role of these loci in the melanic pathway and in building up associations between the coloration and other phenotypic traits through pleiotropy and (iv) discuss how our findings could be used to better understand the mechanisms responsible of the maintenance of the color polymorphism in the European barn owl.

## Results & Discussion

### Genomic and phenotypic landscape of the European barn owl (Tyto alba)

We sampled European barn owls from the western Palearctic, whose coloration varies from white in the south (in the Iberian Peninsula (Portugal - PT), Great Britain (GB) and Levant (Israel -IS)) to dark rufous in the north (especially in Denmark (DK)), with populations in-between (Switzerland (CH) and Greece (GR)) displaying a high color diversity (figure 1a). We conducted whole genome sequencing of 75 individuals from these populations, yielding to a total of 5,112,936 SNPs after filtering. Neutral PCA supported genetic differentiation of the populations (see also [32]), with the first axis separating the Levant lineage (IS) from the rest of the populations, and the second axis isolating the Iberian individuals (PT) samples from the rest of the European populations (Figure S2).

**Figure 1.**
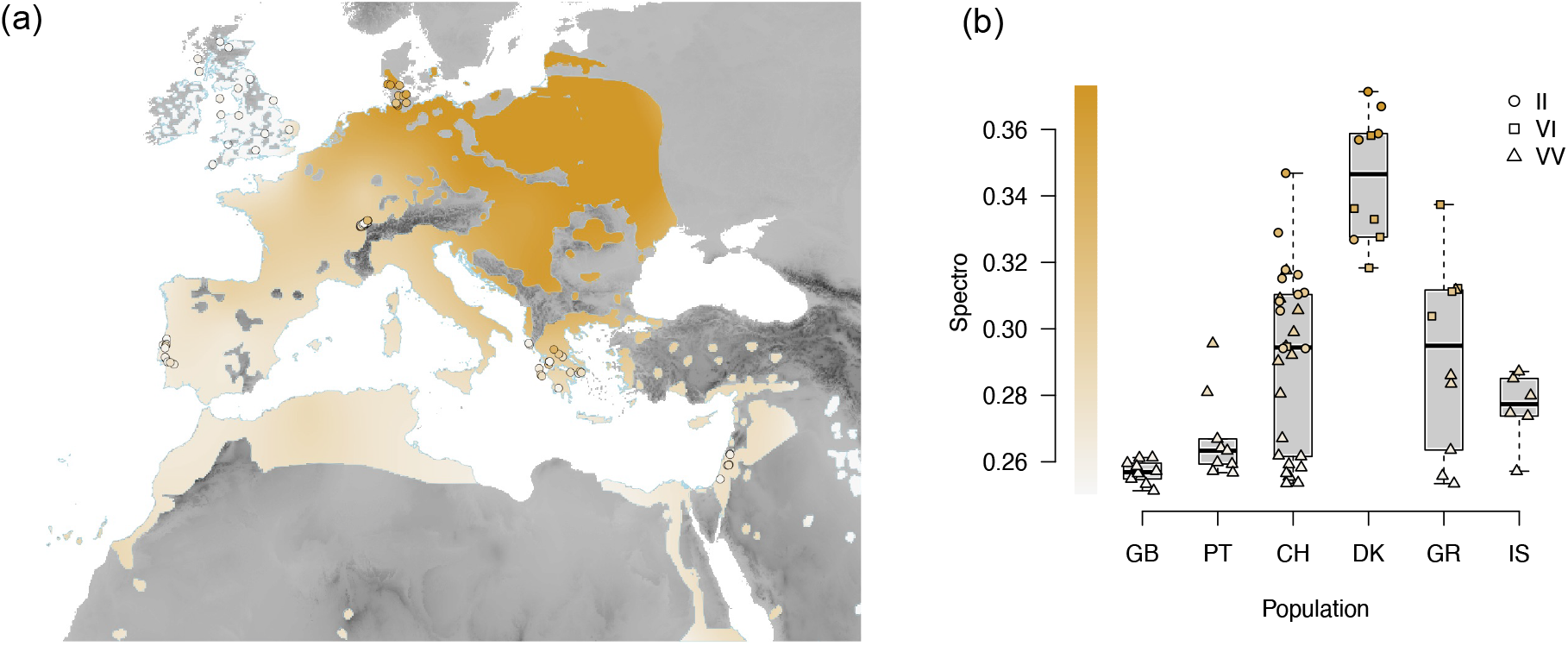
Distribution of the colour phenotype of the barn owl in the western Palearctic. (*a*) the color gradient (interpolated using the Kriging algorithm) displaying gradient of color from white to rufous barn owl in the Western Palearctic. Each dot on the map correspond to one individual, coloured accordingly to its own color. (*b*) Boxplot presenting the variation of the color of the individuals in the different populations sampled. Each dot correspond to one individual, with the shape matching the MC1R genotype and coloured accordingly to its own color.

*MC1R* genotype of these samples was consistent with the color of the individuals, with solely the Valine allele (V) in population with a white phenotype, and the Isoleucine allele (I) present in all populations with rufous individuals, and an increased frequency in the northern population, with only *MC1R*_*VI*_ or *MC1R*_*VV*_ individuals in DK (figure 1b). These results are also in line with the known repartition of this allele at the European scale [19].

### Genome Wide Association identifies a new autosomal region associated with the rufous color

We used a Genome Wide Association (GWA) approach to identify SNPs associated with the coloration of barn owls (Figure 2). Overall, the inflation factor was low (1.05) and most of the points aligned along the 1:1 line in the qq-plot (Figure S4), meaning that the inclusion of the relatedness matrix in the model allowed us to control for population stratification. This GWA identified two outlying SNPs at the genome-wide significance level.

**Figure 2.**
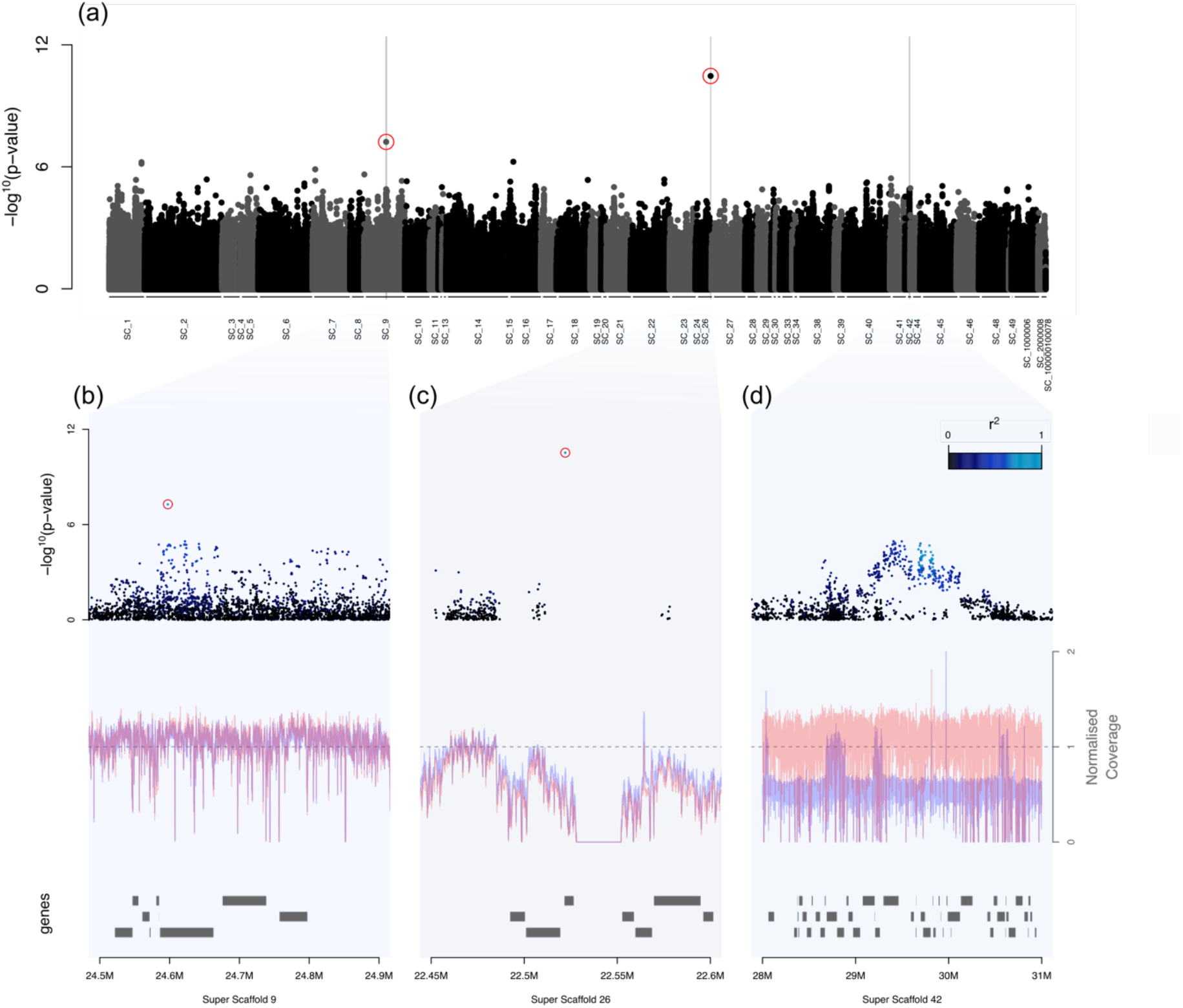
Genome Wide Association study between the genotypes of the Barn Owl and their colour. (*a*) -log^10^(p-values) from the primary test of association with color for each SNPs along the genome. Alternation of colors depict the successive Super-Scaffols (SC_). The two red circles surround significant SNPs presenting a deviation from the 1:1 line of the qq-plot (Sup Fig XXX). Grey bars highlight regions of interest presented in panels (b), (c) and (d). (*b, c, d*) Zooms on different region of the genome displaying a strong signal of association with the color. Dots represent the association test for each SNP in the region of interest. Color of the dots depict their association with the focal SNP within each region (respectively the *MATN2 variant*, the *MC1R variant* and the *Z variant*). Red circles surround the two outlier SNPs in the GWAS. Red and blue line represent the mean normalized coverage for males and females respectively, while the dashed line represent the expected normalized coverage. Rectangles bellow represent the genes annotated in the different regions.

The highest one was located at the *MC1R* locus (hereafter *MC1R variant*, G->A, location: Super-Scaffold 26, pos 22,522,039), with an association score of 5.703e-11 (fig 2a and 2c, value below the Bonferroni’s significance threshold: 0.05 * 5112936 tests = 9.779117e-09). This variant was the one at the *MC1R* gene that we previously discovered using a candidate gene approach [29]. The signal in the region was however surprising. Indeed, if the variant itself showed a high association with the color, the signal was not expanded by linkage disequilibrium to the surrounding variants, all showing a relatively low association with the coloration. This might be due to the low R^2^ with the SNP of interest (Figure 2c) as well as the lower coverage in the region (see coverage fluctuation in panel 2c) probably influenced by the high GC content of the regions [29], making the region poorly sequenced, which may also explain the low number and sparse repartition of the SNPs in the region.

The second highly associated variant was present at the *MATN2* gene (hereafter, *MATN2 variant*, A->G, location: Super-Scaffold 9, pos 24,597,481). It had an association score of 8.813e-08 (fig 2a and 2b) above the Bonferroni’s significance threshold (9.779117e-09) and it could be potentially a type 2 error. However, this SNPs clearly deviated from the 1:1 line of the qq-plot (Figure S4). Moreover, the randomization tests supported that our GWA was not prone to type 2 error signals, with the absence of association as strong as the one observed with the *MATN2* variant in any bootstrap (Figure S5, see *Contribution of multiple loci to the color polymorphism* section for details). The strongest signal in the region came from an intronic region of the *MATN2* gene, but also a cluster of SNPs in linkage disequilibrium, showing a tendency to be associated with the color of the barn owl. We also observed a (non-significant) signal of differentiation of multiple SNPs near the *MTDH* gene, located 160kbp downstream the *MATN2* variant.

### Stratified design in the Swiss population pinpoints a region on the Z chromosome

Previous studies suggested a major role of *MC1R* in the color determinism, and particularly that the allele inducing a whiter plumage coloration (Valine allele - *MC1R-white*) permits the expression of further genetic variation for coloration contrarily to the alternative allele (Isoleucine allele - *MC1R-rufous*), which may epistatically mask the effect of other genetic variation [37]. Because such an expected epistatic effect can hinder QTL discovery, we narrowed down our analysis to consider only the 30 Swiss individuals. This population was chosen as it is one of the populations with the highest variation in coloration (Figure 1b) and there is no apparent genetic structure at the whole genome scale that can be associated with coloration or the *MC1R* variant [32]. Indeed, a neutral PCA shows no differentiation according to coloration nor *MC1R* genotype within the Swiss population (Figure S3). Whole genome F_IS_ for the Swiss population is 0.005, and whole genome F_ST_ between *MC1R*_*VV*_ and *MC1R*_*II*_ in Switzerland is 0.0002, which is consistent with an absence of substructure.

Because of the smaller sample size, we based this GWA on F_ST_ scans rather than a mixed model approach as conducted above for the complete set of samples. We first scanned for highly differentiated genomic regions between the whiter (BC<.28 – n = 10) and more Rufous (BC>.28 – n = 18) phenotypes without considering the MC1R genotype. This first scan showed no strong signal of differentiation along the genome (Figure S6).

We then focused on the 20 Swiss barn owls carrying only the *MC1R-white* allele (i.e MC1R_VV_ individuals), as this variant should permit other genetic variants to have measurable effects on plumage coloration [37]. We scanned for highly differentiated genomic regions between the whiter (BC<.28 – n = 10) and more rufous (BC>.28 – n = 8) *MC1R*_*VV*_ owls, which revealed one highly differentiated region (F_ST_>0.8) harbored on the sex chromosome (Figures S7a and S7b). The association of sex-linked variants to plumage coloration is expected given previous quantitative genetic studies allocating substantial color variation to sex chromosomes in this population [30].

For downstream analyses, we selected the top variant from this region as representative for the genotype at this locus (hereafter called *Z variant*, G->A, location: Super-Scaffold 42, pos 29,829,678). In the GWA, this SNP is non-significant but a cluster of SNPs in linkage disequilibrium in this region, shows a tendency to be associated with the color (Fig 1d)). This region includes multiple genes (Table S2), and among them CHRBP (LOC104362934), located 68,665 bp downstream from the *Z variant*.

### Contribution of multiple loci to the color polymorphism

In order to estimate the contribution of each of the three *loci* identified above (*MATN2 variant*, the *MC1R variant* and the *Z variant*) to barn owl coloration, we fitted an animal model allowing to also estimate the fraction of additive genetic variance that remains unexplained (Figure 3b). The *MC1R locus* had the largest effect on coloration (proportion of variance explained: 0.69, 95% credible interval, CrI: 0.42 -0. 0.93), in line with previous studies focused on a large sampling of Swiss individuals [29,31]. The *Z* locus had a smaller, yet non-negligible effect on coloration (0.09, 95% CrI: 0.03-0.17), while the *MATN2* locus had a small effect with the lower 95% CrI close to zero (0.02, 95% CrI: <0.01-0.06). Moreover, DIC values did not support that including the effect of the *MATN2 locus* has a substantial impact on explaining color variation (ΔDIC = 0.69), contrary to the other *loci* (ΔDIC_*MC1R*_ = 54.66, ΔDIC_*Z* locus_ = 22.03). We thus remain cautious about the role of the *MATN2 locus* in barn owl coloration, despite the clear signal in the GWA. Further investigation including more individuals should allow us to verify the association of the *MATN2 locus* and barn owl coloration.

**Figure 3.**
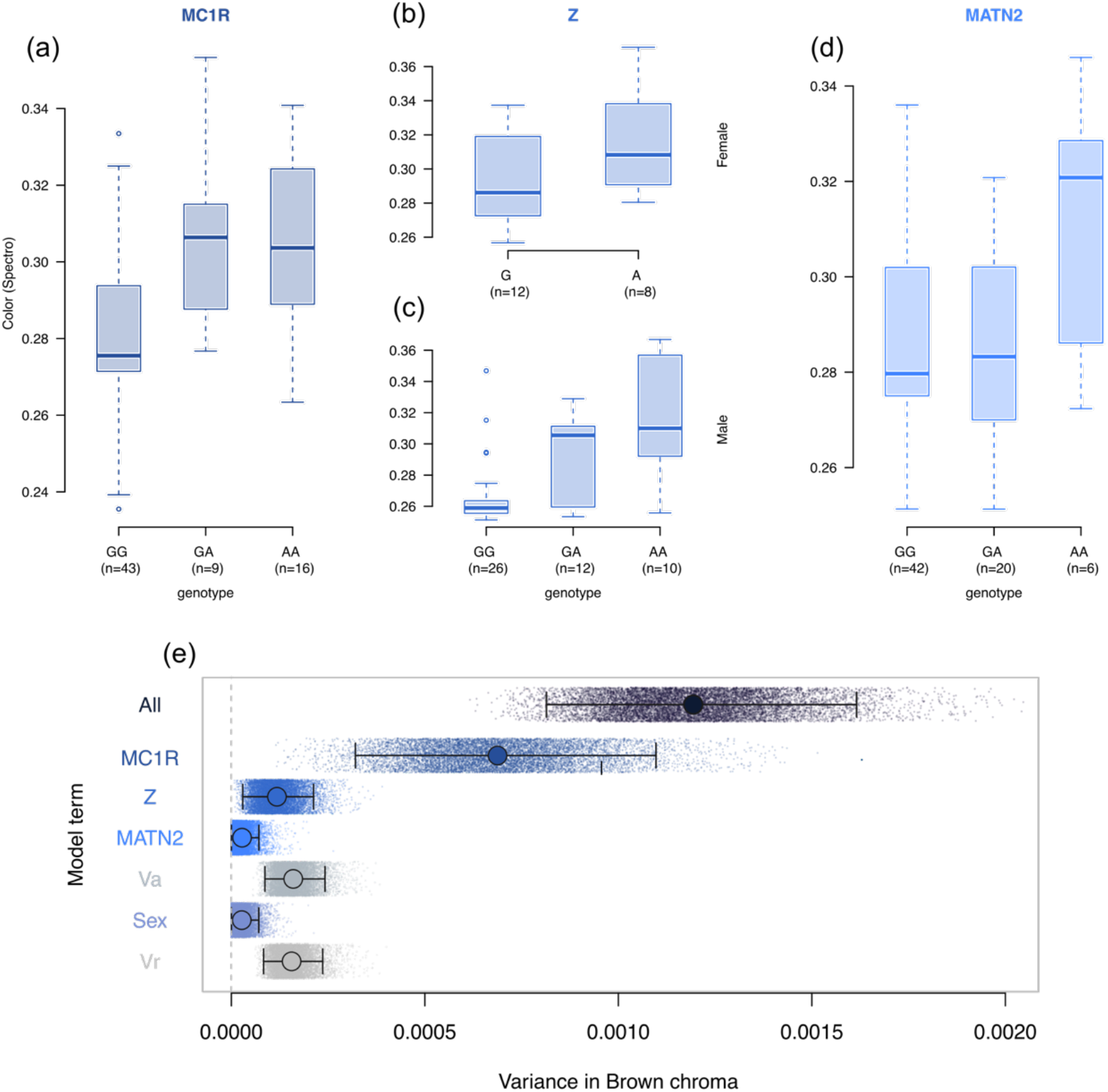
Estimation of the contribution of the different loci to the trait. (*a, b, c, d*) Boxplot of the relation between the genotype at the different locus and the phenotype of the individuals: (*a*) *MC1R variant*, (*b*) *Z variant* for females, (*c*) *Z variant* for males and (*d*) *MATN2 variant*. (*e*) Results of the animal model including the genotype to the 3 loci and the sex of the individuals as fixed predictors. The matrix of relatedness was used as random factor to estimate *Va*. Plain circles depict the variance explained by each term of the model. Credible intervals are presented by the black lines.

We also detected that a non-negligible amount of color variation (0.12, 95% CrI: 0.07-0.19) can still be attributed to genetic variants yet to be discovered. These variants are likely to have a smaller effect on coloration than the variants highlighted in this study. Despite the remaining work to clarify more deeply the genetic architecture sustaining coloration in the barn owl, our study find supports for a rather oligogenic structure with few variants of major effect (particularly the *MC1R* locus).

### Dominance, additive effect, linkage and epistasis

The relation between the genotypes at these three loci and the phenotype (Fig 3a-d) informs us about the dominance interactions between alleles within these loci. For both *MC1R* and *MATN2*, heterozygotes seem to have similar trait value to one of the homozygotes, thus suggesting dominance of one of the alleles over the other. The direction of the dominance is however in opposite direction, with a dominance of the rufous allele at the *MC1R* locus and a dominance of the whitish allele at the *MATN2* locus. On the other hand, the Z locus seems to harbor an additive effect between the two alleles in male diploid individuals at this locus, with heterozygous individuals displaying an intermediate phenotype relatively to the two other homozygotes.

The association between different markers and a trait may be due to physical linkage between them along a chromosome [38]. The three loci we identified are located on independent scaffolds of the assembly of the barn owl (Super-Scaffold 9, 26 and 42 for the *MATN2, MC1R* and the *Z variant* respectively). The location of these regions in other birds’ genomes assembled at chromosome level, shows that they are also located in distinct chromosomes (Table S3). These results, consistent with the known conserved syntheny of bird genomes [39], support an independence of these three markers. The correlation between *MC1R* and *MATN2* may thus be due to selection in the northern environment for the darker rufous phenotype described by Burri et al. [19]. Considering the contribution of the different alleles of the tree markers to the trait (see *Contribution of multiple loci to the color polymorphism* section for details), as well as their different dominance and additive effects within locus, the combined genotype of the individuals at these loci might be sufficient to give a broad panel of color if the effect of the different loci is additive, without taking into account interactions between loci (such as epistasis). The existence of such effects, already hypothesized in the literature [37] could even amplify the number of possible phenotypes. However, the sampling size of this study limits our ability to measure such interactions. Further investigations with an expanded dataset would allow us to measure how these three loci interact to build the red coloration.

### Potential role of newly discovered loci to melanin pathway

Eumelanin and pheomelanin are the two main pigments responsible of variation in coloration in the barn owl [40]. The synthesis of either one or the other of these pigments from the same precursor (tyrosine) relies on a series of reactions that are catalyzed by specific melanogenic enzymes (TYR, TYRP1, TYRP2), regulated by the MITF transcription factor. MITF activity is itself regulated by several signaling pathways that can impact coloration (including MAPK, WNT, PKC, and cAMP) [41]. *MTDH*, a gene located nearby the *MATN2 variant*, has been connected to both MAPK and WNT pathways, notably by down regulating the ERK1/2 signaling [42]. Although the molecular implications of *MTDH* in coloration are still barely understood, variation affecting this gene may interfere with the regulation of MITF, and thus impact the color of the individuals.

Among the pathways regulating MITF, the cAMP pathway is activated by the binding of α-MSH (melanocyte-stimulating hormone) to MC1R, triggering the synthesis of melanin [41]. This gene carrying the non-synonymous *MC1R variant* was already linked to coloration in the barn owl [29] and it has been largely discussed in the literature given its central role in melanin synthesis. The peptidic hormone α-MSH (which binds to MC1R) is produced through the cleavage of the Pro-opiomelanocortin protein, encoded by the *POMC* gene, whose transcription can be activated by the binding of CRH (corticotropin-stimulating hormone) to its receptor CRHR1 [43]. The *Z variant* we found in association with barn owl coloration is in the vicinity of the *CRHBP* gene, which codes the inhibitor of CRH: the CRH binding protein (CRHBP) and may have a potential impact on *POMC* expression and thereby on coloration. The interest of a color variant directly impacting *POMC* expression is that it may also affect the expression of other traits, given the known pleiotropic effects of the melanocortin system [27]. CRH as well as POMC derived hormones participate are main regulators of the stress response [43], which has been previously linked to color variation in the barn owl as well as in other vertebrate species [44]. Thus, further research on the molecular basis of color variation in the barn owl may offer new insights to understand how associations among distinct phenotypes evolves.

### Maintenance of color polymorphism in the European barn owl

Considering the association between polymorphism in melanic coloration and several physiological, behavioral and ecological traits [20-26], combined with its unknow evolutionary origin [32], the question of the mechanisms underlying the maintenance of this polymorphism remains elusive. The three non-mutually exclusive mechanisms proposed so far implies that (i) coloration plays a role in local adaptation at a large European continental scale, favoring darker forms in the north and lighter forms in the south [19,33,34], possibly related to differential foraging strategies and success according to the coloration [22,35]; (ii) coloration is under frequency dependent selection on the different morphs maintain the polymorphism within some populations (i.e. Switzerland or Greece), with darker individuals selected for at low densities and lighter individuals favored at high densities [31]; as well as (iii) sexually antagonist selection favoring darker females (harboring a darker plumage and many black spots located at the tip of ventral body feathers) and lighter males (harboring a lighter plumage and few black spots) in the same polymorphic populations [36]. The knowledge of these new genomic regions associated with the color allows us to establish strategies to further test these hypotheses.

For instance, the local adaptation hypothesis could be tested in the future by measuring traces of selection in these genomic regions. Digging into the specific history of the genomic regions associated with coloration would also allow us to reconstruct its evolutionary history at the continental scale and compare it with the neutral history of the populations. Testing the two other hypotheses (namely the frequency dependent selection and the sexually antagonist selection) will require the combination of both genomic and fitness data. The frequency dependent selection should leave traces at the genomic level. Indeed, since darker individuals seems to be selected for at low densities, rufous variants seem to have a higher fitness the less common they are [31], making it a potential case of negative frequency dependent selection. Such negative frequency dependent selection is thus expected to retain polymorphism in the population [45]. This frequency dependent selection may also be tracked by monitoring the variation of the fitness of the individuals carrying the different alleles according to their frequencies though time.

The sexually antagonist selection hypothesis could also be tested by looking at the correlation of genomic regions associated with the color also with fitness traits depending on the sex of the individual [46]. At the genomic level, such sexually antagonistic selection generates intra-locus sexual conflict that is thought to be resolved through the evolution of sexual chromosomes [47]. This last hypothesis is thus reinforced by the association between a locus on the Z chromosome and the color. Considering that the two coloration extremes are under opposite selection in the two sexes, the fittest males at a given generation (i.e., lighter melanic males, with GG genotype at the *Z variant*) are *de facto* sons of less fit mothers (with a G genotype), while the fittest females (darker) inherited their A allele from a less fit father (either AG or AA).

## Conclusion

This study shed light on the molecular basis of the color polymorphism of the barn owl. By applying methods often limited to model species, we identified three regions acting on the determinism of the plumage coloration of the European barn owl, constituting a first step to understand the molecular basis of this polymorphism. This information helps us to understand the genetic architecture of this trait, giving insight into the potential molecular pathways involved, as well as providing some first clues to disentangle the role played by different forces in maintaining the color polymorphism. Further analyses, to identify causal mutations, explore the history of loci involved in the coloration, as well as their link with evolution through time at population scale, should allow us to better characterize the maintenance of the color polymorphism in the barn owl.

At a phylogeographic scale, several (sub)species of the Tytonidae family exhibit plumage color clines across continents, and it would be worth looking into these loci in other populations and investigate whether these regions are also involved in their color variations. Finally, considering the strong pleiotropy of the melanocortin system, it will be interesting to investigate the potential role on the loci identified in this study on other traits of this fascinating nocturnal raptor. At a broader scale, this study emphasizes the power of quantitative genetics to reveal the molecular basis of polymorphic traits, and thus provide an opportunity to better characterize the relation of the triptych genotype - phenotype - environment, thus building bridges between the ecology and the evolution of species in the wild.

## Material and Methods

### Sampling design, Sequencing and SNPs calling

#### Sampling

In order to cover the phenotypic range of the color of the barn owl in Europe, we retrieved the whole genomes sequences of 55 samples from 6 Western Palearctic localities from Machado et al.2021 and Cumer et al. 2021 (Table S1): 9 individuals from Portugal (PT), 10 from Denmark (DK), 10 from Grand Britain (GB), 10 from Greece (GR) 6 individuals from Israel (IS), and 10 individuals from Switzerland (CH).

The sampling was extended with individuals from Switzerland (CH), a population with a wide range of color variation. On top of the 10 previously described individuals, 20 more were sequenced for this study. These complementary individuals were selected to be mostly homozygous for the MC1R genotype (either VV or II) and preferably males (based on the sexing described in [37]) in order to reduce the effect of differential color between sexes (described by [19]). MC1R_VV_ samples were also selected to cover wide range of color variation. In the final dataset, the Swiss population was represented by 30 individuals including 18 MC1R_VV_ or 11 MC1R_II_ and 1 MC1R_VI_ individuals. Among the MC1R_VV_ 10 were considered as white and 8 rufous (brown chroma of the reflectance spectra <0.28 and >0.28 respectively, see *Phenotypic measurement* section of the Material and Methods for details, Table S1).

#### DNA extraction and Sequencing

For these extra 20 individuals, we followed a similar library preparation and sequencing protocol as described in [48]. In brief, genomic DNA was extracted using the DNeasy Blood & Tissue kit (Qiagen, Hilden, Germany), and individually tagged. 100bp TruSeq DNA PCR-free libraries (Illumina) were prepared according to manufacturer’s instructions. Whole-genome resequencing was performed on multiplexed libraries with Illumina HiSeq 2500 PE high-throughput sequencing at the Lausanne Genomic Technologies Facility (GTF, University of Lausanne, Switzerland).

#### SNPs calling

The bioinformatics pipeline used to obtain analysis-ready SNPs for the dataset including the 75X individuals was adapted from the Genome Analysis Toolkit (GATK) Best Practices [49] to a non-model organism following the developers’ instructions, as in [48]. In brief, raw reads were trimmed with Trimommatic v.0.36 [50] and aligned to the reference barn owl genome [51] with BWA-MEM v.0.7.15 [52]. Base quality score recalibration (BQSR) was performed using high-confidence calls obtained in [32] and following the procedure described in [51].

Genotype calls were then filtered for analyses using a hard-filtering approach as proposed for non-model organisms, using GATK and VCFtools v0.1.14 [53]. Calls were removed if they presented: low individual quality per depth (QD < 5), extreme coverage (800 > DP > 2000) or mapping quality (MQ < 40 and MQ > 70), extreme hetero or homozygosity (ExcessHet > 20 and InbreedingCoeff > 0.9) and high read strand bias (FS > 60 and SOR > 3). We then removed calls for which up to 5% of genotypes had low quality (GQ < 20) and extreme coverage (GenDP < 10 and GenDP > 40). We then filtered to retain only bi-allelic loci, yielding a dataset of 10608379 SNPs. For downstream analyses, SNPs were finally filtered for a minor allele frequency higher or equal to 0.05, yielding to a final dataset of 5,112,936 SNPs.

### Phenotypic information

#### Sex determination based on WGS data

Individual sex was controlled using whole genome information. Mean SNP coverage for autosome (Super Scaffold 1) and Z chromosome (Super Scaffold 42) [51] were extracted using vcftools v0.1.14 [53]. Comparison of both mean coverages allowed to identify two distinct group of individuals, with a ratio close to one for male and 0.5 for female (Figure S1, individual sex based on WGS is reported in Table S1)

#### Phenotypic measurement

Pheomelanin-based color, homogenous on barn owl breast feathers, was measured as the brown chroma of the reflectance spectra (see [33] and [48] for details). Briefly, the brown chroma represents the ratio of the red part of the spectrum (600–700 nm) to the complete visible spectrum (300–700 nm), with higher values indicating larger amounts of reddish pigments on the feathers. The reflectance of four points of the top of three overlapping breast feathers was measured using a S2000 spectrophotometer (Ocean Optics) and a dual deuterium and halogen 2000 light source (Mikropackan, Mikropack). An individual’s brown chroma score was obtained as the average of these points. This method correlates with observational assessments using colour chips (r = –0.78, p < .0001) [54] and has high repeatability (97.6%) [33].

### Neutral diversity, population structure and phenotypic distribution

#### GRM, Kinship Matrix and PCA

Individual-based relatedness (β) [55] and inbreeding coefficient was calculated with the package *SNPRelate* [56] in R (v4.2.2, [57]), with all 75 individuals. In order to avoid redundant signal from linked SNPs, rare (MAF<0.05) alleles were discarded and we trimmed the dataset to only retain SNPs with a r2 lower than 0.4, computed at 500kb max, using the *LD*.*thin()* function from the *gaston* package [58], yielding to a total of 1’033’866 SNPs. The kinship matrix was then transformed into GRM using the *kinship2grm()* function from *hierfstat* package [59]. Two Principal Component Analyses (PCA), including either all individuals or only the 30 Swiss individuals, were also performed with the package *SNPRelate* [56] on the same datasets.

### Identification of genomic regions associated with the color

#### GWAS on European samples

To test for association between genotypes and the color of the 75 European individuals, we used the average information restricted maximum likelihood (AI-REML) algorithm, implemented in the *association*.*test()* function in the *gaston* package [58]. The model included the sex of the individuals as covariable, the Genetic Relationship Matrix (GRM) as random effect, accounting for population structure and cryptic relatedness. We used the Wald test to assess the strength of the association between SNPs and phenotypes. P-values of the test were then compared with a p-values of 0.05 adjusted according to Bonferroni [60] and we visually assessed the deviation of the SNPs from the 1:1 line on a qq-plot.

To validate the association between the color and the genotypes, we repeated the GWA while randomly shuffling phenotypes between individuals. If the GWA has identified real QTL, we would not expect SNPs in randomized tests to deviate from the 1:1 line nor exceed the p-values of 0.05 adjusted according to Bonferroni threshold set above. Across our ten randomized analyses, we did not find any SNP above the Bonferroni threshold nor deviating above the 1:1 plot (Figure S5).

#### Contrast in the Swiss population

In order to detect other loci involved in the coloration of the barn owl, we ran pairwise FST using the *snpgdsFst()* function in *SNPRelate* package [56]. Scans contrasted (i) rufous (i.e. Spectro > 0.28; n=20) and white (i.e. Spectro < 0.28; n= 10) individuals in the 30 swiss panel and (ii) rufous (i.e. Spectro > 0.28; n= 8) and white (i.e. Spectro < 0.28; n= 10) MC1R_VV_ Swiss individuals.

### Syntheny between Assemblies

To measure the potential linkage between the markers associated with the color in the previous sections, we looked at the position of the genes surrounding the variants in different bird genomes assembled at chromosome level, namely the chicken (*Gallus gallus*, GCA_000002315.5, [61]), the collared flycatcher (*Ficedula albicollis*, GCA_000247815.2, [62]) and the golden eagle (*Aquila chrysaetos chrysaetos*, GCA_900496995.4, [63]). Results are presented in the Table S3.

### Variance partition among the color QTLs

To estimate the part of variation in coloration associated to the different *loci* identified during previous steps as well as the remaining unexplained additive genetic variance (*Va*), we fitted an animal model using the *R* package *MCMCglmm* (version 2.34, *R* version 4.1.1) [64]. In this model, we included the fixed effect of the genotype of individuals at the three different *loci* as dosage and sex as fixed predictors. The same matrix of relatedness as the one used in the GWAS was fed to the models to estimate *Va*. Models ran for 103000 iterations, with a burn-in of 3000 and a thinning interval of 10 (effective sampling was ≥ 9384 for all model terms). We calculated the proportion of variance (and the associated 95% credible intervals) as the mean of the posterior distribution of each term (including the residuals, *Vr*) relative to the sum of the posterior distribution of all terms (i.e., the total phenotypic variance).

## Acknowledgements

This work was supported by the Swiss National Science Foundation with grants 31003A-138180 & 31003A_179358 to JG and 31003A_173178 to AR.

## Data Accessibility

The raw Illumina reads for the whole-genome sequenced individuals are available in BioProject PRJNA700797, BioProject PRJNA727977 and BioProject PRJNA925445.

## Author Contribution

TC, APM, LSJ, AR, JG designed this study; LSJ produced whole-genome resequencing libraries with the help of CS; APM mapped the reads and called the variants; TC conducted the analyses with the help of LSJ and suggestions from JG; TC led the writing of the manuscript with input from JG and all co-authors.

## Notes

### Competing Interest Statement

The authors have declared no competing interest.

