## Supplementary material for "The genomic architecture of a continuous color polymorphism in the European barn owl (*Tyto alba*)"

*Supplementary Materials*


SAMPLING DESIGN, SEQUENCING AND SNPS CALLING 2

**Table S1** **2**

PHENOTYPIC INFORMATION 2

**Figure S1** **2**

NEUTRAL DIVERSITY, POPULATIONS STRUCTURE AND PHENOTYPIC DISTRIBUTION 3

**Figure S2** **3**

**Figure S3** **4**

IDENTIFICATION OF GENOMIC REGIONS ASSOCIATED WITH THE COLOR 5

**Figure S4** **5**

**Figure S5** **6**

**Figure S6** **7**

**Figure S7** **7**

**Table S2** **7**

SYNTHENY BETWEEN ASSEMBLIES 8

**Table S3** **8**

### Sampling design, Sequencing and SNPs calling

**Table S1** – Description of the samples used in this study.

### Phenotypic information

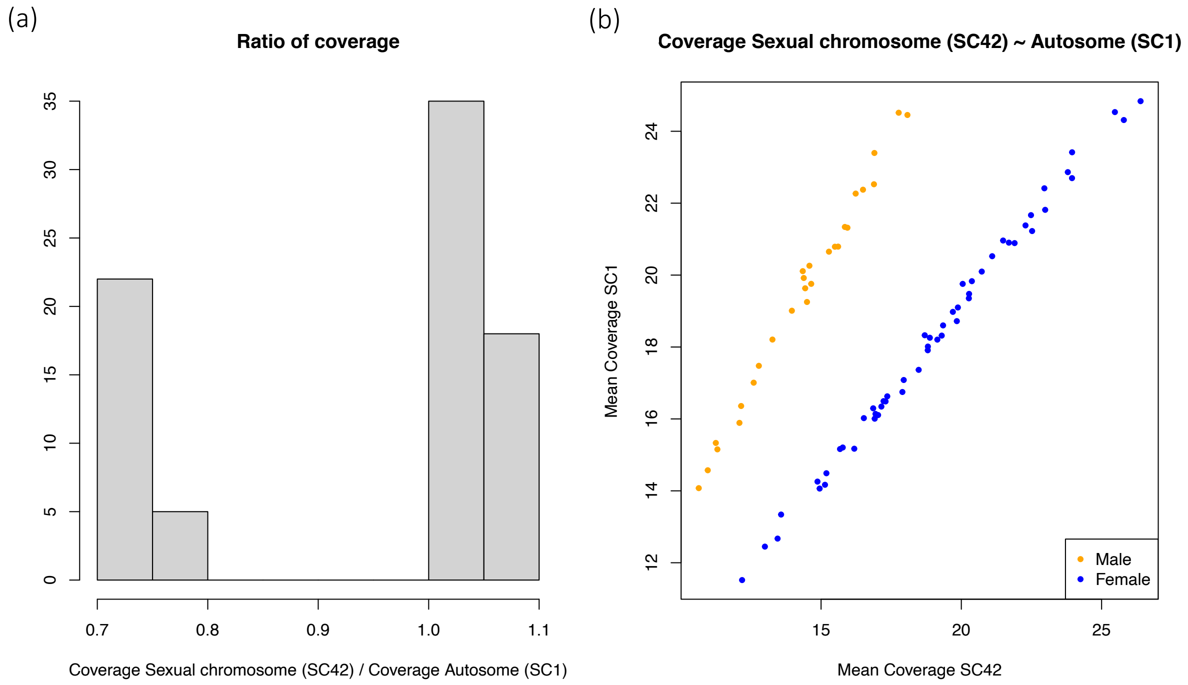


**Figure S1** – Individual sex based on WGS. (*a*) Mean coverages of an autosome (Super Scaffold 1) and the Z chromosome (Super Scaffold 42) allows to identify two distinct group of individuals, with a ratio close and above 1 for males and around 0.7 for females. (*a*) correlation between the mean coverages of an autosome (Super Scaffold 1) and the Z chromosome (Super Scaffold 42) for all individuals. Color is based on the sex attributed based on the ratio of coverage in panel (A), with males in orange and females in blue.

### Neutral diversity, populations structure and phenotypic distribution

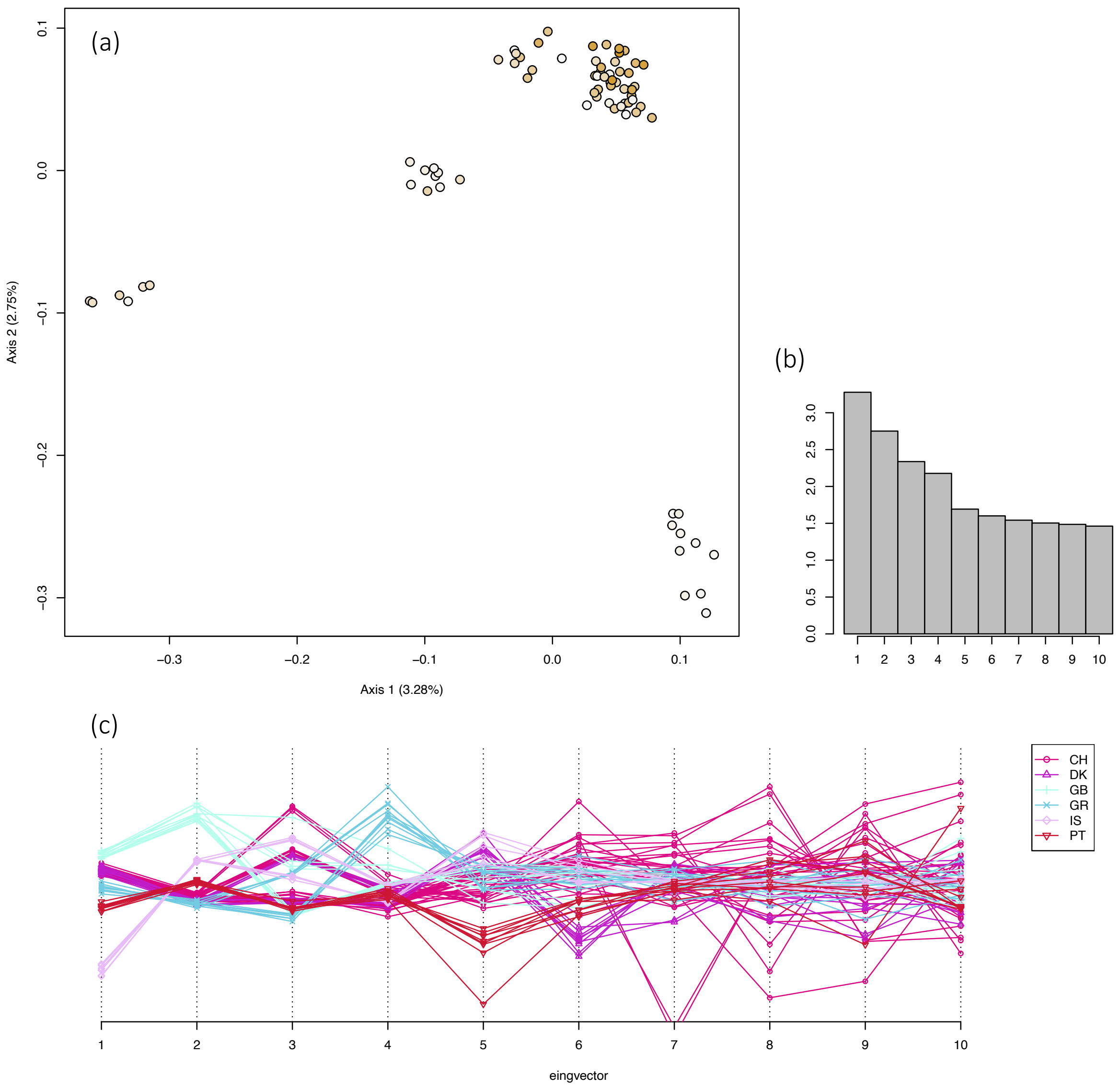


**Figure S2** – Neutral background of the European barn owl (*Tyto alba*). (a) PCA including all European individual, with dots colored according to individual color (scale similar to figure 1). (b) Scree plot of the 10 fist axes of the PCA. (c) Position of the individuals on the 10 first axes of the PCA. Color and type of dot changes accordingly to the population of sampling.

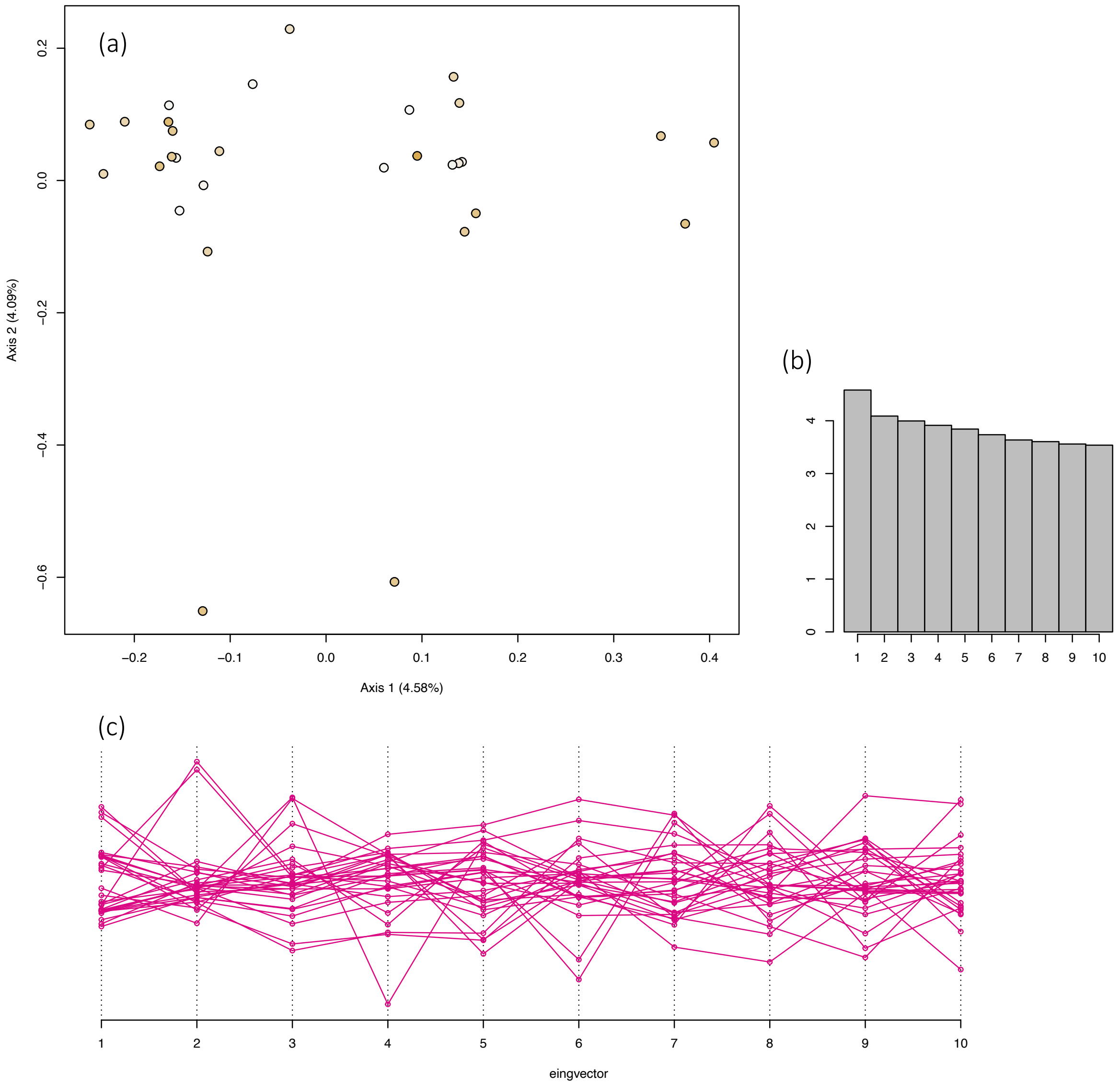


**Figure S3** – Neutral background of Swiss Barn Owl (*Tyto alba*). (a) PCA including the 30 swiss individual, with dots colored according to individual color (scale similar to fig1). (b) Scree plot of the 10 fist axes of the PCA. (c) Position of the Swiss individuals on the 10 first axes of the PCA.

**Identification of genomic regions associated with the color**

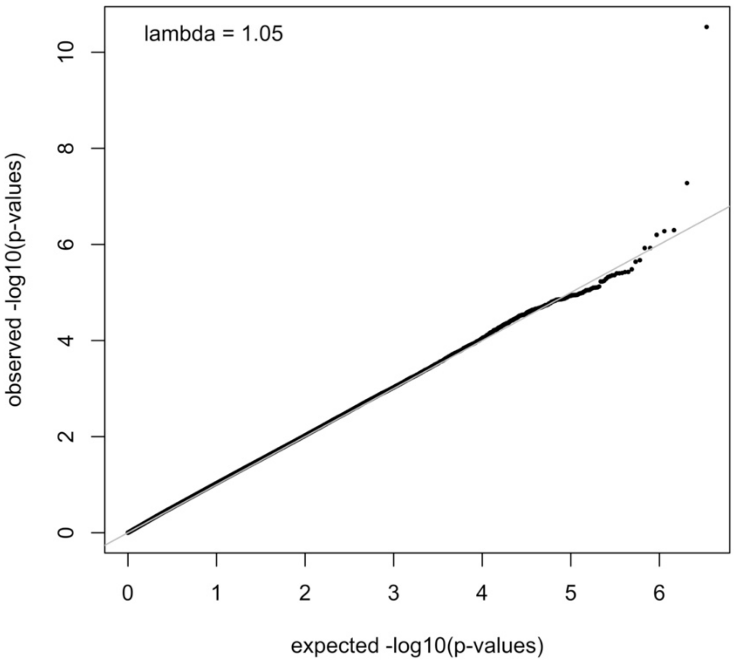


**Figure S4** – QQ-plot of the GWAS presented in Figure 2. Solid grey line represents the 1:1 line.

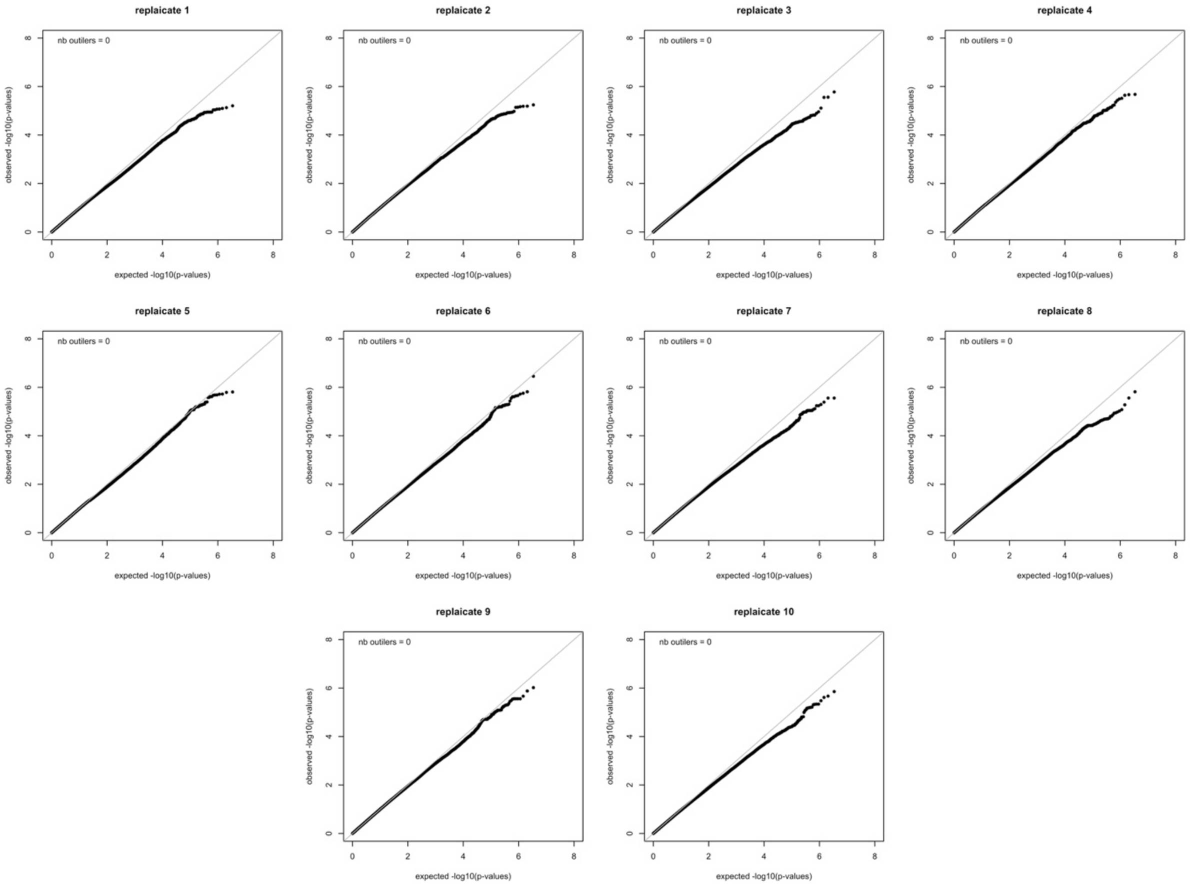


**Figure S5** – QQ-plots for the 10 GWA analyses with randomized phenotype. In each plot, Solid grey line represents the 1:1 line. Top left value confirm that no outliers were detected in the ten replicates according to the Bonferroni corrected p-value.

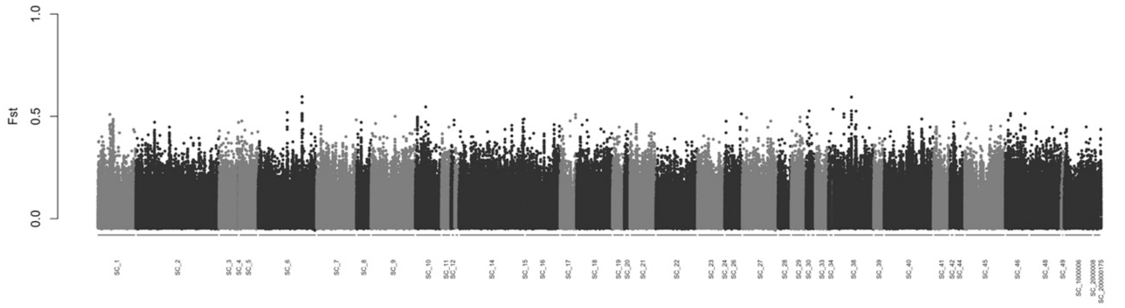


**Figure S6** – Pairwise $F_{ST}$ for each SNP between 10 pale (Spectro<0.28) and 20 rufous (Spectro>0.28) Swiss individuals without accounting for the MC1R genotype.

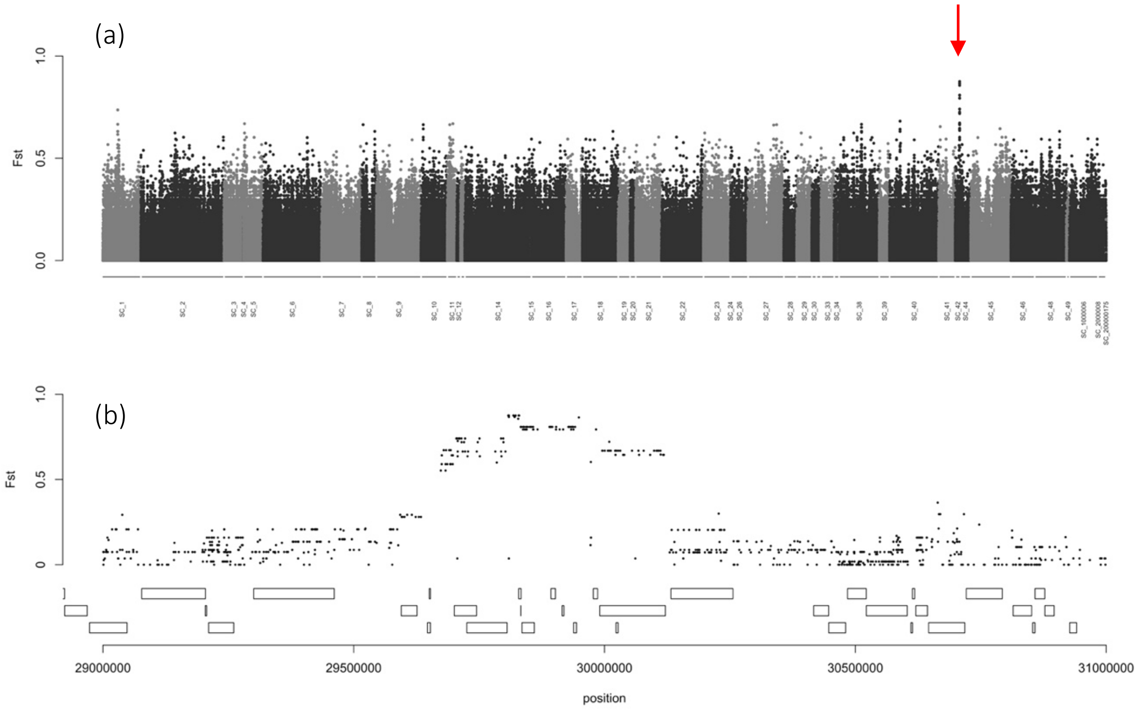


**Figure S7** – (*a*) Pairwise $F_{ST}$ for each SNP between 10 $MC1R_{VV}$ pale (Spectro<0.28) and 8 $MC1R_{VV}$ rufous (Spectro>0.28) swiss individuals. The red arrow point to the region of higher differentiation. (*b*) Zoom on the region on Super-Scaffold 42 showing the highest pic of differentiation. Rectangles bellow represent the genes annotated in the region.

**Table S2** – list of the genes in the regions putatively linked with the color polymorphism in the European Barn Owl.

**Syntheny between Assemblies**

**Table S3** – Chromosomal location of the closest genes of each of the three variants identifies in this study in the genome of the barn owl, the chicken, the flycatcher and the golden eagle.
